# Development and Evaluation of RT-LAMP Assays to Identify Variants of SARS-CoV-2

**DOI:** 10.1101/2022.06.16.496383

**Authors:** Gun-Soo Park, Seong-Jun Kim, Jin-Soo Maeng

## Abstract

Emergence of new variants of Severe Acute Respiratory Syndrome Coronavirus 2 (SARS-CoV-2) during current pandemic of Coronavirus Disease 2019 (COVID-19) and several waves of infections by some of variants emphasized the importance of continuous surveillance. While genomic surveillance through whole genome sequencing is performed as a standard method, identification of known variants through mutation-targeting molecular diagnosis such as qRT-PCR is also useful for timely investigation. However, there are limited studies regarding the concurrent detection and identification of SARS-CoV-2 variants through a LAMP-based method. In this study, we developed and evaluated RT-LAMP assays to detect characteristic deletions of SARS-CoV-2 variants. In addition, we evaluated a fluorescent probe mediated method for identification of single nucleotide substitution by RT-LAMP. Finally, we discussed restrictions and perspectives regarding pathogen screening and surveillance of variants by RT-LAMP based on our observations.

## Introduction

During the COVD-19 pandemic, the evolution of SARS-CoV-2 give rise to many variants. Some of variants are designated as Variants of Concern (VOCs) or Variants of Interest (VOIs) due to known or potential threats to public health (https://www.who.int/en/activities/tracking-SARS-CoV-2-variants/, last accessed March 28, 2022). The risk factors behind the public health threats are higher transmissibility, atypical symptoms, diagnostic failure, decreased effects of therapeutics, and immune evasion to vaccine or previous infections.^1^ VOCs and VOIs are named by Greek alphabet and five variants are designated as VOCS so far: Alpha (Pango lineage^2^ B.1.1.7), Beta (B.1.351), Gamma (P.1), Delta (B.1.617.2) and Omicron (B.1.1.529). These VOCs induced several new waves of epidemics due to its higher transmissibility and/or immune evasion ability.^3-5^ Especially, Omicron is divided into two major sublineages BA.1 and BA.2 (https://github.com/cov-lineages/pango-designation/issues/361), and BA.2 sublineage can outcompete BA.1 sublineage.^6,7^

As evidenced by recurrent emergence of SARS-CoV-2 variant and their phenotypic differences, constant surveillance is important for reaction. The standard method of variants surveillance is whole genome sequencing. Amplicon based partial sequencing can be used in limited way to identify known VOCs.^8^ However, sequencing based identification of variants requires intense facilities and takes more turnaround time compare to qRT-PCR. Therefore, results of routine screening-incorporated qRT-PCR targeting VOC-characterizing mutations can be used as a proxy for some studies such as the timely prevalence of VOCs which are known to be circulating.^9-11^ In fact, assays for various mutations of SARS-CoV-2 including single nucleotide substitutions are readily available.^12^

The gold-standard of COVID-19 diagnosis is qRT-PCR as the method is sensitive, specific and provide quantitative results. qRT-PCR is usually performed in centralized laboratories so that the method has some limitations regarding surveillance such as difficulty to expand test capacity. Therefore, point-of-care test (POCT) oriented methods like antigen-detecting rapid diagnostic tests (RDTs) or isothermal nucleic acid amplification tests (NAATs) are adopted for screening diagnosis.^13^ Especially, many loop-mediated isothermal amplification (LAMP) or recombinase polymerase amplification (RPA) based methods were developed as isothermal NAATs targeting SARS-CoV-2, some with CRISPR-Cas based detection.^14-19^ Nevertheless, there are limited reports regarding identification of VOCs using isothermal NAATs.

In this study, we developed and evaluated reverse-transcription LAMP (RT-LAMP) methods to discriminate VOCs. We basically aimed to design LAMP reaction of which amplification is occurred variant-specifically because such methods are suitable for resource-limited POCT when combined with a simple readout such as colorimetric detection. In addition, we developed an one-step strand displacement probe (OSD-probe) RT-LAMP to discriminate single nucleotide substitution targeting the N501Y mutation of Spike.^20^

## Results

### Target selection

We firstly aimed to discriminate early three VOCs, Alpha, Beta, and Gamma, from non-VOC lineages as Delta was not emerged at the moment. By coincidence, large deletion (9 bp) corresponding to Orf1a SGF3675-3677del within nsp6 region was found in common for the three VOCs so that we chose the region as a target.^21^ We also targeted Spike HV69-70del which is corresponding to S-gene target failure (STGF) of some qRT-PCR diagnostics and Spike N501Y substitution which is another common mutation among early three VOCs but all of the candidate primer sets showed poor sensitivity or failed to discriminate VOC sequences. For the Delta variant, Spike EF156-157del and R158G corresponding 6 bp deletion was selected as a target. For the Omicron-BA.1 variant, Spike GVY142-144del and Y145D corresponding 9 bp deletion and Spike ins214EPE corresponding insertion and we were able to obtain positive results for primer sets targeting the deletion region. Aligned surrounding sequences of target mutation are shown in Supplementary Figure 1. After screening, chosen primer sets were named after affected amino acids (Supplementary Table 1). All the primer sets targeting deletions showed best results with WarmStart Colorimetric LAMP 2X Master Mix.

### Sensitivity and cross-specificity of RT-LAMP targeting large deletions of VOCs

Limit of detection (LoD) against viral RNAs and cross-reactivity to non-target lineages were evaluated. For SGF primer sets, both composition targeting intact or deleted (designated with “del”) sequences showed similar LoD of 50-100 copies/reaction with stochastic detection of 10 copies/reaction samples (Fig. 1A). To test cross-reactivity of primer sets, we used up to 2.5 × 10^6^ copies/reaction of templates considering ∼10^9^ copies/ml of average viral load of SARS-CoV-2.^22^ SGF primer sets showed no cross-reactivity within tested concentrations (Fig. 1B). Omicron-BA.1 has slightly different deletion near the target of SGF primer sets which includes Orf1a L3674 but excludes F3677 so that G-A mismatch occurs at 3’ −5 residue of SGF-FIPdel primer. As a result, LoD of SGFdel primer set to Omicron-BA.1 was 500-1000 copies/reaction (Fig. 1C). SGF primer set showed no cross-reactivity to Omicron-BA.1 within tested concentrations (Fig. 1D).

**Figure 1.**
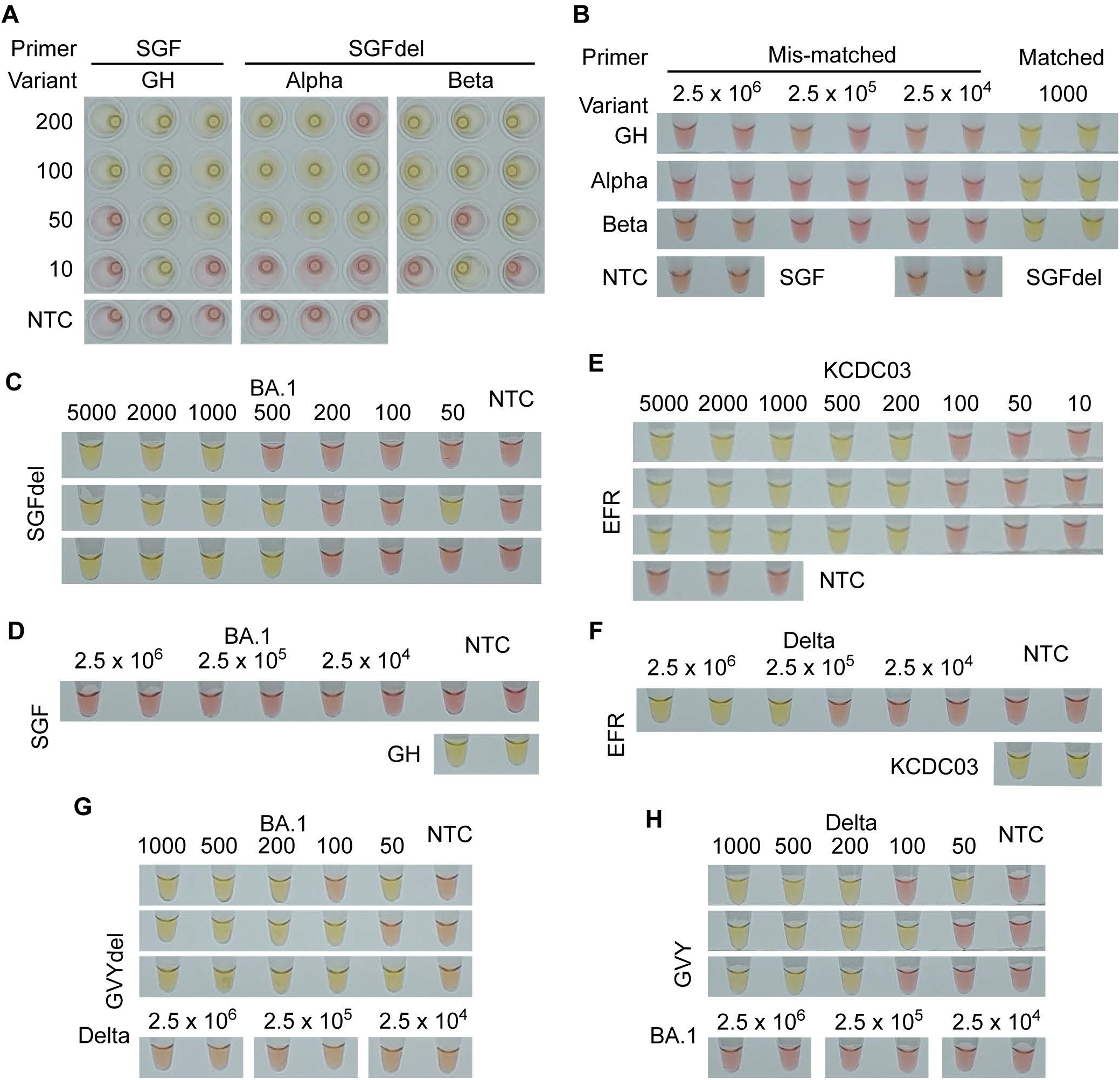
End-point colorimetric results of RT-LAMP targeting deletions. Lineage of viral RNAs, primer sets, and template copy number per reaction are as designated. (A) Triplicate sensitivity test for SGF/SGFdel primer sets. (B) Duplicate cross-reactivity test for SGF/SGFdel primer sets. “Matched” primer-template pairs are of (A) and exchanged for “Mis-matched” samples. (C) Triplicate sensitivity test for SGFdel primer set to Omicron-BA.1 lineage. (D) Duplicate cross-reactivity test for SGF primer set to Omicron-BA.1. 1000 copies/reaction of GH clade viral RNA was used as positive control. (E) Triplicate sensitivity test for EFR primer set. (F) Duplicate cross-reactivity test for EFR primer set. 5000 copies/reaction of wild-type viral RNA (KCDC03) was used as positive control. (G-H) Sensitivity and cross-reactivity test of (G) GVYdel and (H) GVY primer sets. NTC, no template control.

For EFR primer set, non-Delta specific primer set showed positive results during primary screening. With this primer set, therefore, discrimination of Delta variant could be done by target gene amplification failure. LoD of EFR primer set to non-Delta variants is 200 copies/reaction (Fig. 1E). EFR primer set showed cross-reactivity to high copy Delta template from 2.5 × 10^5^ copies/reaction (Fig. 1F).

GVYdel primer set specifically amplify Omicron-BA.1 template with LoD of 50-100 copies/reaction (Fig. 1G). GVY primer set which can amplify non-BA.1 template showed LoD of 200 copies/reaction with stochastic detection of 100 or 50 copies/reaction samples (Fig. 1H). Both GVYdel and GVY primer sets showed no cross-reactivity within tested concentrations. Notably, GVYdel showed late threshold time (Tt) so that incubation time was extended to 90 minutes (Supplementary Fig. 2).

### OSD-probe RT-LAMP targeting Spike N501Y mutation

Although VOCs coincidently have characteristic large deletions which are suitable for discriminating RT-LAMP target, such mutations occur less frequently than single nucleotide substitution.^23^ In addition, single nucleotide substitution can be both characteristic and functionally important.^24,25^ Therefore, we sought to develop a RT-LAMP based diagnostic method to discriminate single nucleotide substitution of SARS-CoV-2. We utilized OSD-probe because the method was reported to distinguish single nucleotide substitution.^20^ OSD-probe functions independently of LAMP primers so that non-specific detection of primer-induced amplification can be avoided and LAMP primer design is less restricted compare to some other fluorescence based detection methods for LAMP.^26^ As a target, Spike N501Y mutation was selected from its convergent emergence in early three VOCs.

OSD-probes did not work properly in WarmStart Colorimetric LAMP 2X Master Mix during primary screen unlike deletion-targeting RT-LAMP. Therefore, *Bst* 2.0 based reaction was adapted. After primer screen, OSD-probe RT-LAMP reaction was optimized for temperature, Betaine usage, concentrations of dNTP and Mg^2+^ ion, and concentration of OSD-probe. While LAMP reaction itself showed LoD of 500-1000 copies/reaction, proper discrimination of mutation can be from 1000-2000 copies/reaction of template (Fig.2A). The requirement of higher concentration of templates for mutation discrimination by OSD-probe is due to its mechanism. OSD-probe can discriminate single nucleotide substitution via different chance of toehold binding and subsequent exchange of strand annealed to fluorescent dye tagged oligonucleotide. Therefore, high concentration of LAMP product is required to observe differential signal by template-probe combinations. To obtain such amount of LAMP product, early beginning of amplification from high concentration of starting template is desired. As a supporting evidence, rather clear difference of end-point fluorescence signals are observed when high-copy templates are used (Fig.2B).

**Figure 2.**
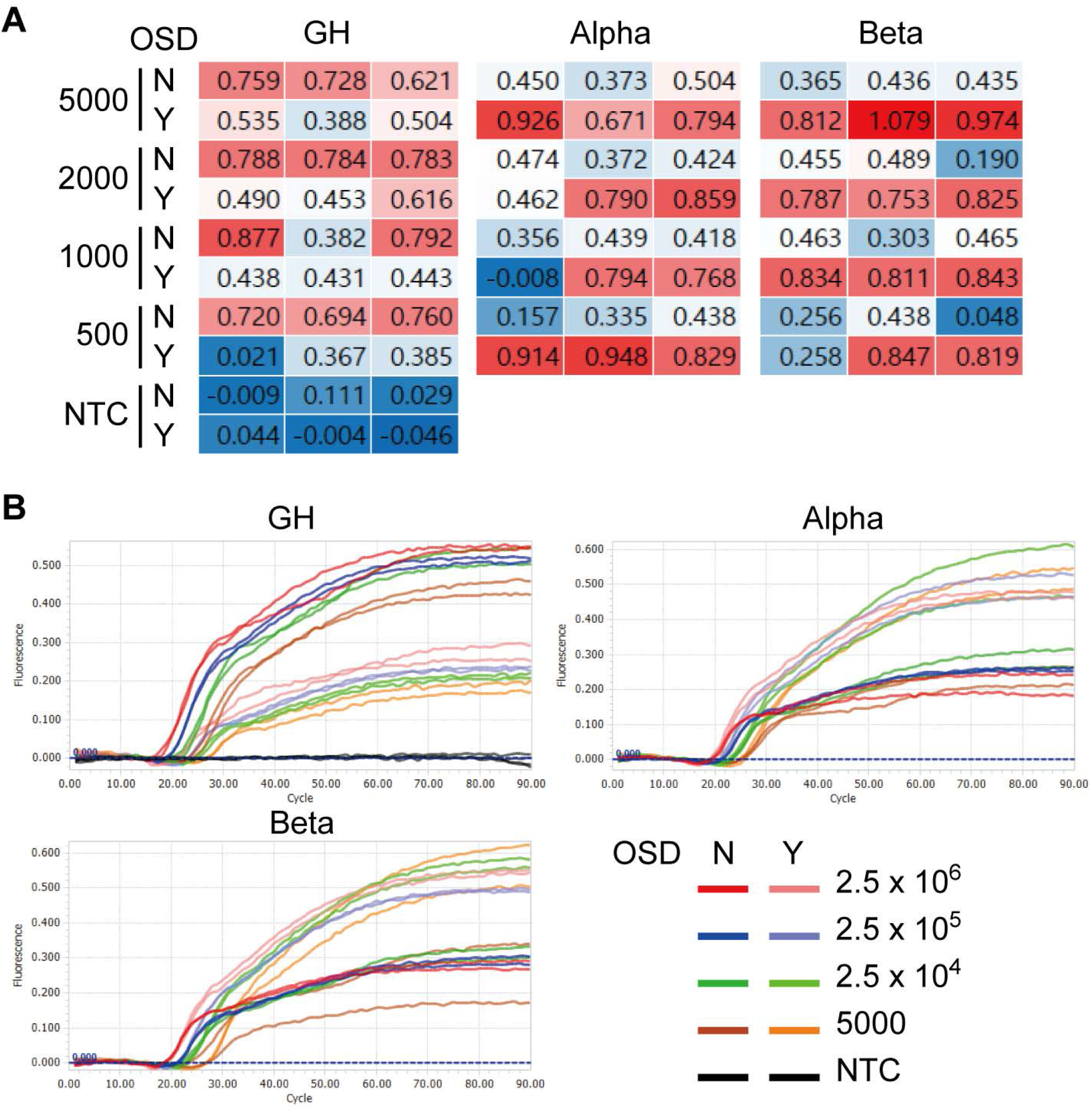
OSD-probe RT-LAMP results. Lineage of viral RNAs, template copy number per reaction, and OSD probe targets are as designated. (A) Base-line corrected end-point fluorescent signal intensities are shown as heat-map for a low-range copy number test. The heatmap was generated by Microsoft Excel program. (B) Real-time FAM fluorescence signal of OSD probes for a high-range copy number test. NTC, no template control.

As Omicron variants emerged, we tested OSD-probe RT-LAMP for Spike N501Y with BA.1 viral RNA. As a result, template induced LAMP reaction was only observed for reactions with high RNA copies. OSD-probe signal was not properly generated, consistent with varying Tt of LAMP reaction (Supplementary Fig.3). The reason would be many mutations in F1/B1 primer region; three residues in F1 region of Omicron, three or two residues in B1 region of BA.1 or BA.2, respectively, are mutated verses each primer sequences.

## Discussion

As a new variant of pathogen emerges during an epidemic, the variant may be detected by genomic surveillance or epidemiologic surveillance followed by confirmation by whole genome sequencing. Then, the degree of threats to public health by new variants should be quickly assessed by their epidemiologic features such as reproduction number or by phenotypic assays regarding immune evasion and response to treatments.^27^ Upon the results of aforementioned assays, the variant may be declared as a VOI or a VOC then public health policies may be changed including active monitoring of the variant. The monitoring includes identification of the variant either by sequencing or by molecular diagnosis targeting characteristic mutations. Because emergence and spread of such variants can be rapid as witted from cases of SARS-CoV-2^5^, concurrent detection of pathogen and identification of a certain variant by molecular diagnostic methods is favorable than sequencing due to different turnaround time. Additionally, a method for the active monitoring should be both sensitive to fulfill diagnostic purpose and rapidly adaptable to new mutations for a case that the target variant is characterized by a novel mutation. While qRT-PCR meets the criteria, partially due to controlled primer-probe annealing through thermal cycling, adaptability of isothermal NAATs requires further evaluations. For this purpose, we developed and evaluated RT-LAMP methods to identify SARS-CoV-2 VOCs. Especially, we designed RT-LAMP to amplify templates from specific target variants to keep resource-limited POCT oriented virtues of LAMP.

To summarize, we were able to develop RT-LAMP methods to discriminate VOCs by targeting large deletions. By targeting Orf1a SGF3675-3677del of Alpha, Beta, Gamma and Omicron-BA.2, LoD of 50-100 copies/reaction was observed with no cross-reactivity up to 2.5 × 10^6^ copies/reaction. For Omicron-BA.1, ten-fold increased LoD was observed due to different deletion position result in single nucleotide mismatch in a LAMP primer. By targeting Delta-specific Spike EF156-157del and R158G corresponding deletion, LoD of 200 copies/reaction was observed for non-Delta variant with no cross reactivity up to 2.5 × 10^4^ copies/reaction of Delta template. For Omicron-BA.1 specific Spike GVY142-144del and Y145D corresponding deletion, LoD of 50-200 copies/reaction was observed with no cross reactivity up to 2.5 × 10^6^ copies/reaction. Additionally, we introduced an OSD-probe RT-LAMP method to discriminate Spike N501Y mutation.

We partially succeeded in development of concurrent RT-LAMP methods for COVID-19 screening and VOCs identification. First, RT-LAMP for simple binary readouts such as colorimetric detection was possible for codon-scale mutations. As we screened LAMP primers for both criteria of sensitivity and VOC specificity, our failure of designing RT-LAMP primer for Spike N501Y mutation does not negate potential of LAMP based detection of single nucleotide mutations. In fact, allele-specific LAMP methods were possible for various targets.^28,29^ Nevertheless, mutation detection by LAMP primers only may not be applied in general cases. For example, EFR primer set showed cross-reactivity from 2.5 × 10^5^ copies/reaction of Delta template. This cross-reactivity would be from sequence similarity at direct downstream of the deletion region (AGTTcA – gAGTTtA, Supplementary Fig.1C). Due to relatively poor sensitivity of LAMP based methods compare to PCR based methods, efforts to increase LAMP sensitivity were made.^30-33^ Considering the mechanism of LAMP amplification which rely on thermodynamic chance of primer invasion and loop formation, such efforts seem to result in increased robustness and decreased specificity contrarily. On the other hand, adapting touchdown procedure for semi-controlled primer binding in LAMP increased both sensitivity and specificity.^34^

To accomplish both of sensitive screening and specific identification through LAMP, detection of mutations can be performed by sequence specific methods independent of LAMP reaction.^29^ In this case, LAMP primer can be optimized as long as the target mutation reside within a amplicon. Simple oligonucleotide probes such as OSD-probe or molecular beacon are reported to be able to discriminate single nucleotide variation with LAMP.^20,35^ Both methods are successfully applied for multiplexed or variant detection of SARS-CoV-2 with LAMP.^36,37^ Nevertheless, OSD-probe and molecular beacon require fluorescence detection as a readout and targeting loop region of LAMP product so that the design of LAMP primers is still somewhat restricted. CRISPR-Cas based detection methods are of particular interest because the system can detect single nucleotide variations, is multiplexable and have versatile readout methods.^38,39^ Additionally, detection target can be any region within the amplicon for widely used Cas12 or Cas13 methods; Cas12 can target both single-stranded or double-stranded DNA and RNA generation step is incorporated for Cas13.^40,41^ Indeed, studies detecting SARS-CoV-2 mutations through CRISPR-Cas systems were reported.^42-44^

Another obstacle for SARS-CoV-2 variants identification is multiplexing. Since many mutations are convergently emerged in subsets of variants, combinations of mutations should be determined for identification of a certain variant. From the POCT point of view, using multiple fluorescent dyes is not favorable because accompanying instrument would become expensive. One option is lateral flow assay (LFA). Simple readout provided by LFA is beneficial for POCT especially when it is performed by minimally trained personnel and multiplexed test is possible by using differently labelled primers. Therefore, LFA have been utilized as a LAMP readout.^45-47^ While coupling of LAMP and LFA requires well-designed mutation specific LAMP reactions for variant identification, such LAMP can be hard to develop as we discussed. At this point, detection of mutations by CRISPR-Cas system would be applicable because; (1) LFA, LAMP and CRISPR-Cas detection can be merged^17^, and (2) CRISPR-Cas based multiplexing and single nucleotide variation is possible.^38,44^ Another option is to run multiple reactions by targets using multi-compartment design.^48-51^ Multi-compartment design may require more amount of sample and reagent compare to single-tube multiplexing which would be coupled with LFA. However, cross-contamination would be minimized as no extra step is required after amplification.

From the diagnostic point of view, the priority is to avoid detection failure since the identification of variants is not a primary goal of diagnosis.^52^ With such priority, LAMP has intrinsic weakness for concurrent pathogen detection and variant identification regardless of selection of detection methods. LAMP reaction requires up to eight primer binding sites, including loop primers, aligned to form characteristic dumbbell structure within around 200 bp target region. Each primer binding sites should have desirable sequences with criteria like proper GC percentage, no long repeat of single base and minimum primer-primer interaction. Naturally, limited regions on a genome would be proper target of LAMP and indeed selected, partially due to limited tools for primer design.^15^ Unless target mutations for variant identification overlap a proper LAMP amplicon region, LoD of the mutation targeting LAMP would be higher compare to detection-oriented LAMP. Indeed, LoD of some primer sets used in this study – EFR, GVY and S501 – were higher than previously reported SARS-CoV-2 detecting LAMP primers.^14,17,53^

Constant monitoring and alarming of new variants is especially important for hypothetical origin of VOCs/VOIs such as a chronic patient, wild animals, or patients in a place lacking sufficient genomic surveillance capacity.^27^ Diagnosis-coupled detection of new mutation would be effective as target subjects requires routine tests or tests should be done in place-of-care. For such monitoring, the large coverage of target amplicon by LAMP primers would be beneficial. When a mutation occurs within LAMP primer binding sites, sensitivity and threshold time would be affected. The most effective targets would be mutation hot-spots such as receptor binding motif of SARS-CoV-2. Indeed, the sensitivity of S501 primer set was affected by mutations introduced in Omicron (Supplementary Fig.3). Otherwise, sequencing can be adapted taking advantage of relatively longer amplicon size of LAMP than qPCR. For instance, a place-of-care oriented method utilizing nanopore sequencing was reported.^54^ Finally, pathogen detection-oriented LAMP should be included in the test to avoid missing infections and as a test-positivity comparing control. The target should be both conserved and abundant. In case of SARS-CoV-2, a conserved region in *N*-gene would serve the purpose as the target template copy number is highest among Orfs by the composition of sub-genomic RNAs.^55-57^ In conclusion, we developed and evaluated RT-LAMP methods to detect and discriminate SARS-CoV-2 VOCs. Based on our observations, restrictions and potentials of LAMP based methods are discussed in the perspective of reacting to pathogen’s variants.

## Methods

### Viral RNA

SARS-CoV-2 viral RNA of wild-type lineage (BetaCoV/Korea/KCDC03/2020, provided by Korea Disease Control and Prevention Agency) and Delta variant (NCCP 43390, provided by Korea National Culture Collection for Pathogens) were prepared as previously described.^58^ Viral RNA from GH clade (NCCP 43345), Alpha variant (NCCP 43381), Beta variant (NCCP 43382), and Omicron-BA.1 variant (NCCP 43408) were provided by Korea National Culture Collection for Pathogens.

The copy number of viral RNAs were titrated by qRT-PCR using Luna Universal Probe One-Step RT-qPCR Kit (NEB) and LightCycler 96 instrument (Roche). Primers and probes are listed in Supplementary Table 2. *In vitro* transcribed standard RNAs were prepared from cloned fragments containing each amplicon of qRT-PCR. Briefly, RNAs were synthesized with MEGAscript T7 Transcription Kit (Thermo Scientific) and purified with Zymoclean Gel RNA Recovery Kit (Zymo Research) after electrophoresis using native 1x MOPS buffered agarose gel.

### RT-LAMP

LAMP primers were designed based on suggestions given by PrimerExplorer V5 (http://primerexplorer.jp/lampv5e/index.html, last accessed March 28, 2022) for each target. The working concentration of LAMP primers are as follow: 1.6 μM for inner primers (FIP/BIP), 0.2 μM for outer primers (F3/B3), and 0.4 μM for loop primers (LF/LB). All RT-LAMP reactions were performed with 2 μl of templates in TE buffer (10 mM Tris-Cl, pH 7.5, and 1 mM EDTA) for total 15 μl reaction volume. For deletion-targeting RT-LAMP, WarmStart Colorimetric LAMP 2X Master Mix (NEB) was used with 0.4 μM SYTO-9 (Thermo Scientific).

For OSD-probe RT-LAMP, OSD-probes were designed and prepared following instructions in previous reports.^20,36^ Working concentration of OSD-probe was 50 nM by fluorescent probe and probe to quencher ratio was 1:5. Other components of OSD-probe RT-LAMP than template, primers and probe are as follow: 1x Isothermal Amplification Buffer I [NEB; 20 mM Tris-HCl, pH 8.8, 10 mM (NH4)2SO4, 50 mM KCl, 2 mM MgSO4, and 0.1% Tween 20], 4 mM MgSO4 (NEB; final, 6 mM Mg^2+^), 1 mM each dNTP (Enzynomics), 0.4 μM SYTO-82 (Thermo Scientific), 6 U *Bst* 2.0 WarmStart DNA polymerase (NEB), and 2.25 U WarmStart® RTx Reverse Transcriptase (NEB). Leuco crystal violet solution was added for end-point colorimetric detection as previously described.^14^ RT-LAMP reactions were performed using LightCycler 96 instrument and fluorescence signals were measured for every minute. For deletion-targeting RT-LAMP, reactions were performed for 40 minutes at 65°C. For OSD-probe LAMP, reactions were performed for 90 minutes at 60°C. Any changed conditions are specified.

## Supplementary Information

Supplementary tables, figures and their legends are provided as supplementary information.

## Acknowledgements

Authors thank to Korea Centers for Disease Control and Prevention (KCDC) for kind and rapid sharing of isolated strain of SARS-CoV-2. This work was supported by the National Research Council of Science and Technology (NST) grant by the Ministry of Science and ICT (Grant No. CRC-16-01-KRICT and GN160600-KFRI).

## Author Contributions

G.-S.P. designed and performed experiments. S.-J.K. provided samples. G.-S.P. and J.-S.M. wrote the manuscript.

## Competing Interests

The authors declare no competing interests.

## Data Availability

No datasets were generated or analyzed during the current study.

## Notes

### Competing Interest Statement

The authors have declared no competing interest.

## References

1 World Health Organization. Guidance for surveillance of SARS-CoV-2 variants: Interim guidance, 9 August 2021. (2021).

2 Rambaut, A. et al. A dynamic nomenclature proposal for SARS-CoV-2 lineages to assist genomic epidemiology. Nat Microbiol 5, 1403–1407, doi:10.1038/s41564-020-0770-5 (2020).

3 Tao, K. et al. The biological and clinical significance of emerging SARS-CoV-2 variants. Nat Rev Genet 22, 757–773, doi:10.1038/s41576-021-00408-x (2021).

4 Viana, R. et al. Rapid epidemic expansion of the SARS-CoV-2 Omicron variant in southern Africa. Nature 603, 679–686, doi:10.1038/s41586-022-04411-y (2022).

5 World Health Organization. Enhancing readiness for omicron (B. 1.1. 529): technical brief and priority actions for member states. (2022).

6 Lyngse, F. P. et al. Transmission of SARS-CoV-2 Omicron VOC subvariants BA.1 and BA.2: Evidence from Danish Households. medRxiv, 2022.2001.2028.22270044, doi:10.1101/2022.01.28.22270044 (2022).

7 Iketani, S. et al. Antibody evasion properties of SARS-CoV-2 Omicron sublineages. Nature, doi:10.1038/s41586-022-04594-4 (2022).

8 European Centre for Disease Prevention and Control & World Health Organization Regional Office for Europe. Methods for the detection and characterisation of SARS-CoV-2 variants – first update. (2021).

9 Davies, N. G. et al. Estimated transmissibility and impact of SARS-CoV-2 lineage B.1.1.7 in England. Science 372, doi:10.1126/science.abg3055 (2021).

10 Wolter, N. et al. Early assessment of the clinical severity of the SARS-CoV-2 omicron variant in South Africa: a data linkage study. Lancet 399, 437–446, doi:10.1016/S0140-6736(22)00017-4 (2022).

11 Lentini, A., Pereira, A., Winqvist, O. & Reinius, B. Monitoring of the SARS-CoV-2 Omicron BA.1/BA.2 variant transition in the Swedish population reveals higher viral quantity in BA.2 cases. medRxiv, 2022.2003.2026.22272984, doi:10.1101/2022.03.26.22272984 (2022).

12 Lai, E. et al. A method for variant agnostic detection of SARS-CoV-2, rapid monitoring of circulating variants, detection of mutations of biological significance, and early detection of emergent variants such as Omicron. medRxiv, 2022.2001.2008.22268865, doi:10.1101/2022.01.08.22268865 (2022).

13 World Health Organization. Public health surveillance for COVID-19: Interim guidance, 14 February 2022. (2022).

14 Park, G. S. et al. Development of Reverse Transcription Loop-Mediated Isothermal Amplification Assays Targeting Severe Acute Respiratory Syndrome Coronavirus 2 (SARS-CoV-2). J Mol Diagn 22, 729–735, doi:10.1016/j.jmoldx.2020.03.006 (2020).

15 Alves, P. A. et al. Optimization and Clinical Validation of Colorimetric Reverse Transcription Loop-Mediated Isothermal Amplification, a Fast, Highly Sensitive and Specific COVID-19 Molecular Diagnostic Tool That Is Robust to Detect SARS-CoV-2 Variants of Concern. Front Microbiol 12, 713713, doi:10.3389/fmicb.2021.713713 (2021).

16 Dong, Y. et al. Comparative evaluation of 19 reverse transcription loop-mediated isothermal amplification assays for detection of SARS-CoV-2. Sci Rep 11, 2936, doi:10.1038/s41598-020-80314-0 (2021).

17 Broughton, J. P. et al. CRISPR-Cas12-based detection of SARS-CoV-2. Nat Biotechnol 38, 870–874, doi:10.1038/s41587-020-0513-4 (2020).

18 Ding, X. et al. Ultrasensitive and visual detection of SARS-CoV-2 using all-in-one dual CRISPR-Cas12a assay. Nat Commun 11, 4711, doi:10.1038/s41467-020-18575-6 (2020).

19 Patchsung, M. et al. Clinical validation of a Cas13-based assay for the detection of SARS-CoV-2 RNA. Nat Biomed Eng, doi:10.1038/s41551-020-00603-x (2020).

20 Jiang, Y. S. et al. Robust strand exchange reactions for the sequence-specific, real-time detection of nucleic acid amplicons. Anal Chem 87, 3314–3320, doi:10.1021/ac504387c (2015).

21 Martin, D. P. et al. The emergence and ongoing convergent evolution of the SARS-CoV-2 N501Y lineages. Cell 184, 5189–5200 e5187, doi:10.1016/j.cell.2021.09.003 (2021).

22 Puhach, O. et al. Infectious viral load in unvaccinated and vaccinated patients infected with SARS-CoV-2 WT, Delta and Omicron. medRxiv, 2022.2001.2010.22269010, doi:10.1101/2022.01.10.22269010 (2022).

23 Sanjuan, R., Nebot, M. R., Chirico, N., Mansky, L. M. & Belshaw, R. Viral mutation rates. J Virol 84, 9733–9748, doi:10.1128/JVI.00694-10 (2010).

24 Korber, B. et al. Tracking Changes in SARS-CoV-2 Spike: Evidence that D614G Increases Infectivity of the COVID-19 Virus. Cell 182, 812–827 e819, doi:10.1016/j.cell.2020.06.043 (2020).

25 Wang, P. et al. Antibody resistance of SARS-CoV-2 variants B.1.351 and B.1.1.7. Nature 593, 130–135, doi:10.1038/s41586-021-03398-2 (2021).

26 Becherer, L. et al. Loop-mediated isothermal amplification (LAMP) – review and classification of methods for sequence-specific detection. Analytical Methods 12, 717–746, doi:10.1039/C9AY02246E (2020).

27 Oude Munnink, B. B. et al. The next phase of SARS-CoV-2 surveillance: real-time molecular epidemiology. Nat Med 27, 1518–1524, doi:10.1038/s41591-021-01472-w (2021).

28 Gill, P. & Hadian Amree, A. AS-LAMP: A New and Alternative Method for Genotyping. Avicenna J Med Biotechnol 12, 2–8 (2020).

29 Varona, M. & Anderson, J. L. Advances in Mutation Detection Using Loop-Mediated Isothermal Amplification. ACS Omega 6, 3463–3469, doi:10.1021/acsomega.0c06093 (2021).

30 Kimura, Y. et al. Optimization of turn-back primers in isothermal amplification. Nucleic Acids Res 39, e59, doi:10.1093/nar/gkr041 (2011).

31 Khorosheva, E. M., Karymov, M. A., Selck, D. A. & Ismagilov, R. F. Lack of correlation between reaction speed and analytical sensitivity in isothermal amplification reveals the value of digital methods for optimization: validation using digital real-time RT-LAMP. Nucleic Acids Res 44, e10, doi:10.1093/nar/gkv877 (2016).

32 Zhou, Y. et al. A Mismatch-Tolerant Reverse Transcription Loop-Mediated Isothermal Amplification Method and Its Application on Simultaneous Detection of All Four Serotype of Dengue Viruses. Front Microbiol 10, 1056, doi:10.3389/fmicb.2019.01056 (2019).

33 Zhang, Y. et al. Enhancing colorimetric loop-mediated isothermal amplification speed and sensitivity with guanidine chloride. Biotechniques 69, 178–185, doi:10.2144/btn-2020-0078 (2020).

34 Wang, D. G., Brewster, J. D., Paul, M. & Tomasula, P. M. Two methods for increased specificity and sensitivity in loop-mediated isothermal amplification. Molecules 20, 6048–6059, doi:10.3390/molecules20046048 (2015).

35 Varona, M. & Anderson, J. L. Visual Detection of Single-Nucleotide Polymorphisms Using Molecular Beacon Loop-Mediated Isothermal Amplification with Centrifuge-Free DNA Extraction. Anal Chem 91, 6991–6995, doi:10.1021/acs.analchem.9b01762 (2019).

36 Bhadra, S., Riedel, T. E., Lakhotia, S., Tran, N. D. & Ellington, A. D. High-Surety Isothermal Amplification and Detection of SARS-CoV-2. mSphere 6, doi:10.1128/mSphere.00911-20 (2021).

37 Sherrill-Mix, S., Van Duyne, G. D. & Bushman, F. D. Molecular Beacons Allow Specific RT-LAMP Detection of B.1.1.7 Variant SARS-CoV-2. J Biomol Tech 32, 98–101, doi:10.7171/jbt.21-3203-004 (2021).

38 Kaminski, M. M., Abudayyeh, O. O., Gootenberg, J. S., Zhang, F. & Collins, J. J. CRISPR-based diagnostics. Nat Biomed Eng 5, 643–656, doi:10.1038/s41551-021-00760-7 (2021).

39 Tang, Y. et al. The CRISPR-Cas toolbox for analytical and diagnostic assay development. Chem Soc Rev 50, 11844–11869, doi:10.1039/d1cs00098e (2021).

40 Chen, J. S. et al. CRISPR-Cas12a target binding unleashes indiscriminate single-stranded DNase activity. Science 360, 436–439, doi:10.1126/science.aar6245 (2018).

41 Schermer, B. et al. Rapid SARS-CoV-2 testing in primary material based on a novel multiplex RT-LAMP assay. PLoS One 15, e0238612, doi:10.1371/journal.pone.0238612 (2020).

42 Patchsung, M. et al. A multiplexed Cas13-based assay with point-of-care attributes for simultaneous COVID-19 diagnosis and variant surveillance. medRxiv, 2022.2003.2017.22272589, doi:10.1101/2022.03.17.22272589 (2022).

43 Nguyen, L. T. et al. A Thermostable Cas12b from *Brevibacillus* Leverages One-pot Detection of SARS-CoV-2 Variants of Concern. medRxiv, 2021.2010.2015.21265066, doi:10.1101/2021.10.15.21265066 (2021).

44 de Puig, H. et al. Minimally instrumented SHERLOCK (miSHERLOCK) for CRISPR-based point-of-care diagnosis of SARS-CoV-2 and emerging variants. Sci Adv 7, doi:10.1126/sciadv.abh2944 (2021).

45 Nurul Najian, A. B., Engku Nur Syafirah, E.A., Ismail, N., Mohamed, M. & Yean, C. Y. Development of multiplex loop mediated isothermal amplification (m-LAMP) label-based gold nanoparticles lateral flow dipstick biosensor for detection of pathogenic Leptospira. Anal Chim Acta 903, 142–148, doi:10.1016/j.aca.2015.11.015 (2016).

46 Zhang, Y., Yu, Y. & Ying, J. Y. Multi-Color Au/Ag Nanoparticles for Multiplexed Lateral Flow Assay Based on Spatial Separation and Color Co-Localization. Advanced Functional Materials 32, 2109553, doi:https://doi.org/10.1002/adfm.202109553 (2022).

47 Zhu, X. et al. Multiplex reverse transcription loop-mediated isothermal amplification combined with nanoparticle-based lateral flow biosensor for the diagnosis of COVID-19. Biosens Bioelectron 166, 112437, doi:10.1016/j.bios.2020.112437 (2020).

48 Poritz, M. A. et al. FilmArray, an automated nested multiplex PCR system for multi-pathogen detection: development and application to respiratory tract infection. PLoS One 6, e26047, doi:10.1371/journal.pone.0026047 (2011).

49 Song, J. et al. Two-Stage Isothermal Enzymatic Amplification for Concurrent Multiplex Molecular Detection. Clin Chem 63, 714–722, doi:10.1373/clinchem.2016.263665 (2017).

50 Zhu, Y. S. et al. Multiplex and visual detection of African Swine Fever Virus (ASFV) based on Hive-Chip and direct loop-mediated isothermal amplification. Anal Chim Acta 1140, 30–40, doi:10.1016/j.aca.2020.10.011 (2020).

51 Nguyen, H. V. et al. Nucleic acid diagnostics on the total integrated lab-on-a-disc for point-of-care testing. Biosens Bioelectron 141, 111466, doi:10.1016/j.bios.2019.111466 (2019).

52 Thomas, E., Delabat, S., Carattini, Y. L. & Andrews, D. M. SARS-CoV-2 and Variant Diagnostic Testing Approaches in the United States. Viruses 13, doi:10.3390/v13122492 (2021).

53 Rabe, B. A. & Cepko, C. SARS-CoV-2 detection using isothermal amplification and a rapid, inexpensive protocol for sample inactivation and purification. Proc Natl Acad Sci U S A 117, 24450–24458, doi:10.1073/pnas.2011221117 (2020).

54 James, P. et al. LamPORE: rapid, accurate and highly scalable molecular screening for SARS-CoV-2 infection, based on nanopore sequencing. medRxiv, 2020.2008.2007.20161737, doi:10.1101/2020.08.07.20161737 (2020).

55 Kim, D. et al. The Architecture of SARS-CoV-2 Transcriptome. Cell 181, 914–921 e910, doi:10.1016/j.cell.2020.04.011 (2020).

56 Doddapaneni, H. et al. Oligonucleotide capture sequencing of the SARS-CoV-2 genome and subgenomic fragments from COVID-19 individuals. PLoS One 16, e0244468, doi:10.1371/journal.pone.0244468 (2021).

57 Ogando, N. S. et al. SARS-coronavirus-2 replication in Vero E6 cells: replication kinetics, rapid adaptation and cytopathology. J Gen Virol 101, 925–940, doi:10.1099/jgv.0.001453 (2020).

58 Jung, Y. et al. Comparative Analysis of Primer-Probe Sets for RT-qPCR of COVID-19 Causative Virus (SARS-CoV-2). ACS Infect Dis 6, 2513–2523, doi:10.1021/acsinfecdis.0c00464 (2020).

