## supplementary information for "Development and Evaluation of RT-LAMP Assays to Identify Variants of SARS-CoV-2"

**Supplementary Table 1.** Sequences of LAMP primers and OSD probes

| Set Name | Primer | Sequence^ |
|---|---|---|
| **SGF** | F3 | 5’ – GCGTATTATGACATGGTTGG – 3’ |
| | B3 | 5’ – ACTTTATAAACGAGTGTCAAGAC – 3’ |
| | FIP | 5’ – ACAGCTGATGCATACATAACACA TATGGTTGATACTAGTTTGTCTGG – 3’ |
| | FIPdel* | 5’ – ACAGCTGATGCATACATAACACA TATGGTTGATACTAGTTT*G*AAGC – 3’ |
| | BIP | 5’ – TAATCCTTATGACAGCAAGAACTGT ATTCATAAGTGTCCACACTCTC – 3’ |
| | LB | 5’ – GTATGATGATGGTGCTAG – 3’ |
| **EFR** | F3 | 5’ – GTGAATTTCAATTTTGTAATGATCC – 3’ |
| | B3 | 5’ – AAACCCTGAGGGAGATCA – 3’ |
| | FIP | 5’ – CCATAAGAAAAGGCTGAGAGACATA GTTGGATGGAAAGTGAGTTCA – 3’ |
| | BIP | 5’ – ACCTTGAAGGAAAACAGGGTAATT AATTAATAGGCGTGTGCTTAG – 3’ |
| | LF | 5’ – CAAAAGTGCAATTATTCGCACTAG – 3’ |
| | LB | 5’ – TCAAAAATCTTAGGGAATTTGTGTT – 3’ |
| **GVY** | F3 | 5’ – CTAATGTTGTTATTAAAGTCTGTGA – 3’ |
| | B3 | 5’ – ATTCTTAAACACAAATTCCCTAAG – 3’ |
| | FIP | 5’ – CACTTTCCATCCAACTTTTGTTG GTAATGATCCATTTTTGGGTGT – 3’ |
| | FIPdel | 5’ – AACTCTGAACTCACTTTCCATCC GTAATGATCCATTTTTGGACCA – 3’ |
| | BIP | 5’ – TCTAGTGCGAATAATTGCACTTTTG ACCCTGTTTTCCTTCAAGGT – 3’ |
| | LB | 5’ – ATGTCTCTCAGCCTTTTCTTATGG – 3’ |
| **S501** | F3 | 5’ – CTGTATAGATTGTTTAGGAAGTCT – 3’ |
| | B3 | 5’ – CTTTTTAGGTCCACAAACAGT – 3’ |
| | FIP | 5’ – TYAACACCATTACAAGGTGTGCT AATCTCAAACCTTTTGAGAGAG – 3’ |
| | BIP | 5’ – TACAATCATATGGTTTCCAACC CTGGTGCATGTAGAAGTTCAA – 3’ |
| | LF | 5’ – CCGGCCTGATAGATTTCAGTTG – 3’ |
| | LB | 5’ – CCAACCATACAGAGTAGTAGTACT – 3’ |
| | Probe_N | 5’ – /6-FAM/ ACTACTACTCTGTATGGTTGGTAACCAACACCATAAGTG /Inverted dT/– 3’ |
| | Probe_Y | 5’ – /6-FAM/ ACTACTACTCTGTATGGTTGGTAACCAACACCATTAGTG /Inverted dT/– 3’ |
| | Quencher | 5’ – GTTGGTTACCAACCATACAGAGTAGTAGT /BHQ1/ – 3’ |

^ Residues affected by mutations are designated by underline.

* Mismatching nucleotide residue to Omicron-BA.1 was designated by *italic*.

**Supplementary Table 2.** Sequences of qRT-PCR primers and probes

| Set Name | Primer | Sequence | Working Concentration (nmol/L) |
|---|---|---|---|
| **Nsp3^** | F | 5' – GAAGAGCAAGAAGAAGATTGGT – 3' | 400 |
| | R | 5' – TGGTGTAAGTTCCATCTCTAATTG – 3' | 400 |
| | P | 5' – /6-FAM/ ACGGCAGTGAGGACAATCAGACA /BHQ1/ – 3' | 200 |
| **S3*** | F | 5' – GGTGATTCTTCTTCAGGTTGGA – 3' | 400 |
| | R | 5' – GTACACTTTGTTTCTGAGAGAGG – 3' | 400 |
| | P | 5' – /6-FAM/ CTGGTGCTGCAGCTTATTATGTGGG /BHQ1/– 3' | 200 |

^ Nsp3 set was used to titrate Orf1 copy number of viral RNAs corresponding to SGF/SGFdel LAMP.

* S3 set was used to titrate Spike copy number of viral RNAs corresponding to EGR/GVY/GVYdel/S501 LAMP.

### *Supplementary Figure 1*

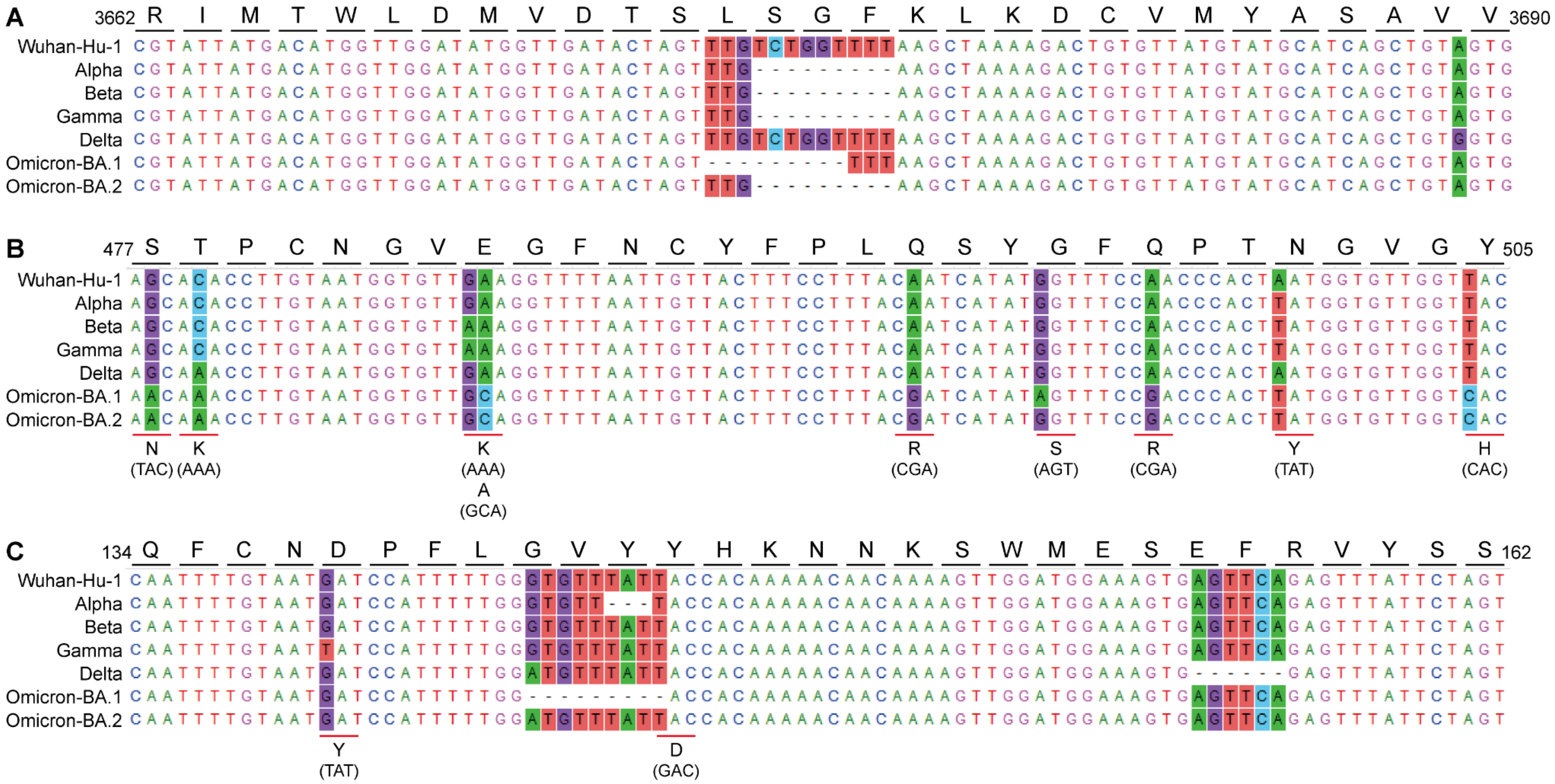


**Supplementary Figure 1.** Aligned sequences (A) around Orf1a SGF3675-3677del mutation, (B) of Spike S477 to Y505 related to N501Y targeting OSD-probe RT-LAMP, and (C) around Spike EF156-157del+R158G and GVY142-144del+Y145D mutations. Corresponding amino acids are designated above of each panel. Changed amino acids by mutations are designated by red lines with corresponding codons. Wuhan-Hu-1 sequence is the NCBI reference sequence (NC_045512.2). Sequences of variants are obtained from GISAID (https://www.gisaid.org/) and accession ID are as follow: Alpha – EPI_ISL_601443, Beta – EPI_ISL_660190, Gamma – EPI_ISL_833137, Delta – EPI_ISL_1409773, Omicron-BA.1 – EPI_ISL_7545782, and Omicron-BA.2 – EPI_ISL_7190366.

### *Supplementary Figure 2*

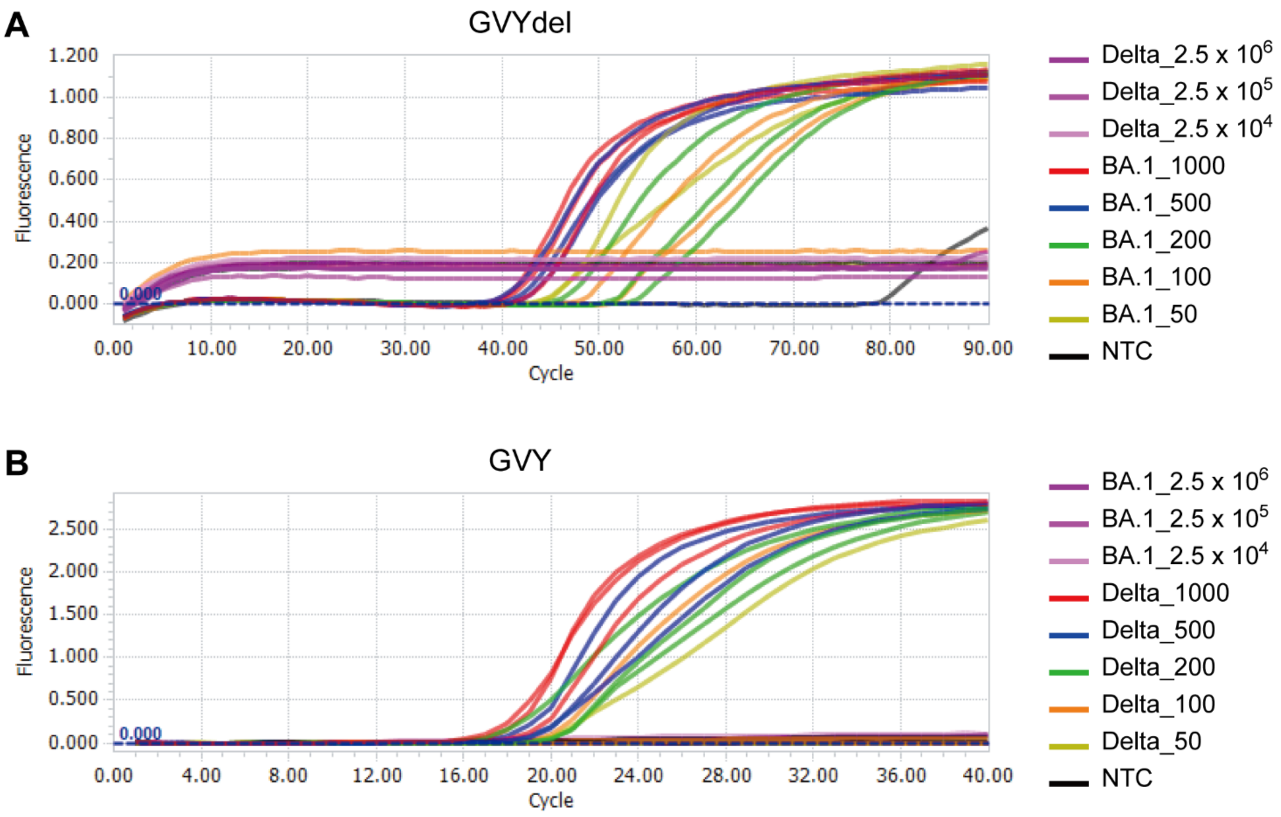


**Supplementary Figure 2.** Real-time SYTO-9 fluorescence signal of GVYdel (A) and GVY (B) primer sets corresponding to panel (G) and (H) of main Figure 1, respectively.

### Supplementary Figure 3

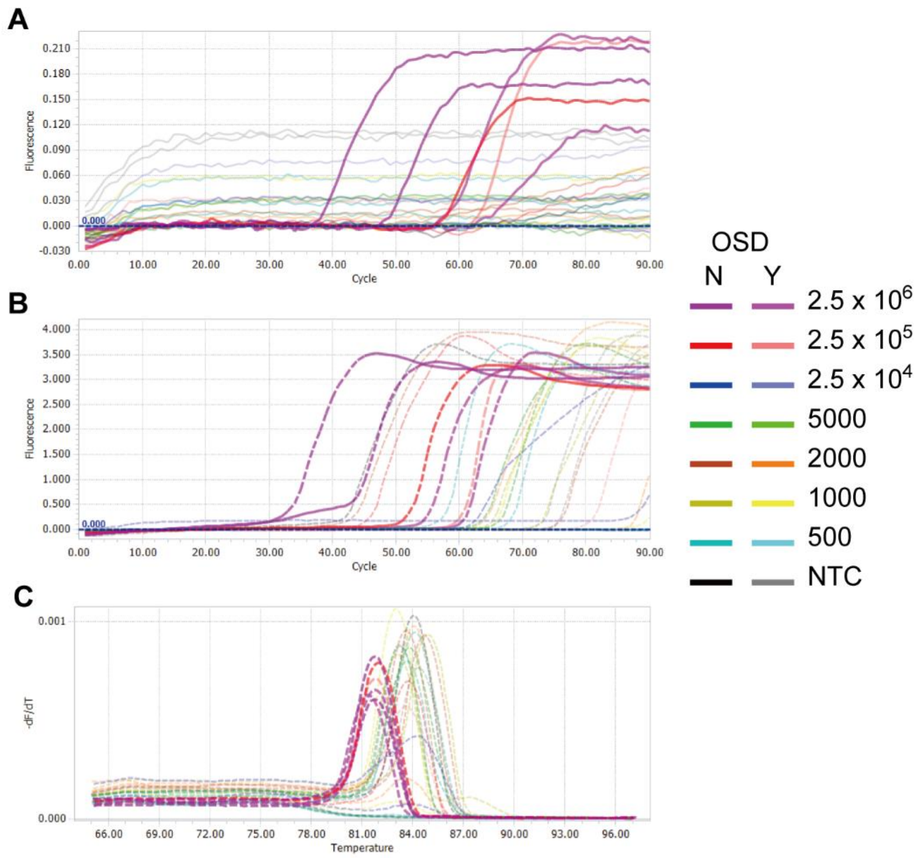


**Supplementary Figure 3.** Real-time fluorescence signal of (A) FAM from OSD-probes, (B) SYTO-82, and (C) melting curve of SYTO-82 from OSD-probe RT-LAMP test with Omicron-BA.1 viral RNA. Copy number per reaction and OSD-probe targets are indicated for each color of lines. Template-derived proper amplifications are indicated by bolded lines.
